# DNA damage-induced phosphorylation of histone H2A at serine 15 is linked to DNA end resection

**DOI:** 10.1101/2021.02.08.430376

**Authors:** Salar Ahmad, Valérie Côté, Jacques Côté

## Abstract

The repair of DNA double-strand breaks (DSBs) occurs in chromatin and several histone post-translational modifications have been implicated in the process. Modifications of histone H2A N-terminal tail has also been linked to DNA damage response, through acetylation or ubiquitination of lysine residues that regulate repair pathway choice. Here, we characterize a new DNA damage-induced phosphorylation on chromatin, at serine 15 of H2A in yeast. We show that this SQ motif functions independently of the classical S129 C-terminal site (γH2A) and mutant mimicking constitutive phosphorylation increases cell sensitivity to DNA damage. H2AS129ph is induced by Tel1^ATM^ and Mec1^ATR^, and loss of Lcd1^ATRIP^ or Mec1 signaling decreases γH2A spreading distal to the DSB. In contrast, H2AS15ph is completely dependent on Lcd1^ATRIP^, indicating that this modification only happens when end resection is engaged. This is supported by an increase of RPA and a decrease in DNA signal near the DSB in the *H2AS-15E* phosphomimic mutant, indicating higher resection. This serine is replaced by a lysine in mammals (H2AK15), which undergoes an acetyl-monoubiquityl switch to regulate binding of 53BP1 and resection. This regulation seems functionally conserved with budding yeast H2AS15 and 53BP1-homolog Rad9, using different post-translational modifications between organisms but achieving the same function.

## INTRODUCTION

The genetic information of the cell is challenged by various endogenous and exogenous cues which leads to DNA damage. If unchecked, this can lead to mutations, translocations, loss of genetic information and, in higher eukaryotes, be the underlying cause for diseases such as cancer (1, 2). One of the most harmful form of DNA damage is DNA double-strand break (DSB). To endure and mitigate such damage to the DNA, cells orchestrate cell cycle arrest and DNA repair pathways, collectively termed the DNA damage response (DDR). Two main pathways to repair DSBs are non-homologous end joining (NHEJ) and homologous recombination (HR). NHEJ involves direct ligation of the two broken DNA ends which may require some processing, making it a more error-prone pathway (3). In HR, single-stranded DNA (ssDNA) is generated through a process called resection and involves the use of homologous sequence or sister chromatid as a template to copy the genetic information, ensuring a lower error rate (4).

The phosphatidylinositol-3-OH kinase-related kinases (PIKKs) family plays an important role in DDR. It includes *S. cerevisiae* Tel1 and Mec1 and mammalian ataxia telangiectasia-mutated (ATM), AT-related (ATR) and DNA-dependent protein kinase (DNA-PK) (5, 6). One of the first DNA damage-dependent chromatin modification to be identified was γ-H2A(X), which is the phosphorylation of histone H2A at S129 in yeast and H2A.X at S139 in mammals (7-9). This phosphorylation is carried out by both Tel1/ATM and Mec1/ATR. In yeast, H2AS129ph spreads more than 50Kb on both sides flanking the DSB, whereas in mammals it covers up to a megabase (10, 11). This histone mark is implicated in the DNA damage response and is required for proper recruitment of several repair/signaling factors as well as chromatin remodelers/modifiers (12-17). H2AX^-/-^ mice are radiation sensitive, growth retarded, immune deficient and show recruitment defect for a variety of repair factors such as Nbs1, 53BP1 and BRCA1 (18).

Interestingly, budding yeast has another Tel1/Mec1-dependent SQ phosphorylation site at H2B C-terminus, H2BT129ph, which largely correlates with H2AS129ph but is modulated by it (10). Other DNA damage-induced phosphorylation marks have been mapped on chromatin but are not deposited by PIKKs. These include H4S1ph, H2AS122ph and H2AT126ph in yeast, which have been linked to processes like chromatin restoration after repair, meiosis, HR and repair of fragile sites with CAG/CTG repeats (19-23). Apart from phosphorylation, other histone modifications like acetylation, methylation, ubiquitylation and SUMOylation have been implicated in DDR (16). In mammals H2AK15 is ubiquitinated by RNF168 and this mark is responsible for recruitment and repair foci formation of 53BP1 in combination with H4K20me2, inhibiting resection and favoring NHEJ (24). At DSBs, the TIP60 complex can acetylate H2AK15 and binds H4K20me, thus antagonizing 53BP1 and favoring repair by HR (25). In budding yeast, Rad9 is the functional homolog of 53BP1, contains a Tudor domain which binds H3K79me (equivalent to mammalian H4K20) and a BRCT domain which recognizes γ-H2A (17, 26, 27). Both domains are required for the recruitment and retention of Rad9 at DSBs (17, 28). In contrast to 53BP1, Rad9 does not have a ubiquitin-binding domain neither the yeast H2A has lysine corresponding to human H2AK15 (**Fig. 1A**). Instead, yeast H2A has a SQ site at this position which has been found phosphorylated in a Mec1-dependent manner upon DNA damage, in a large phosphoproteomic study (29).

**Fig 1:**
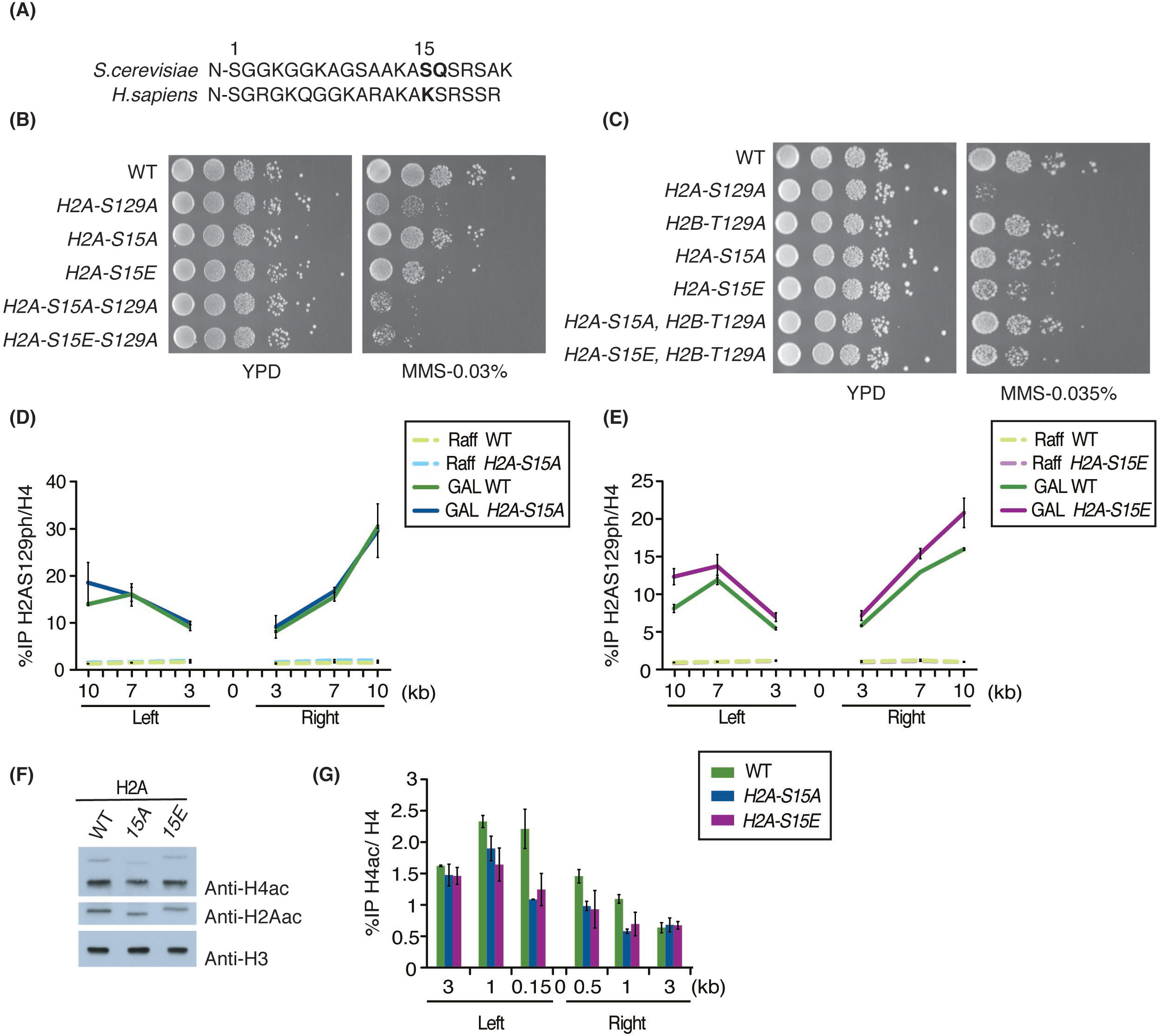
*H2A-S15* mutants are sensitive to DNA damage and functions independently of γ-H2A. (A) Sequence of H2A N-termini highlighting the SQ site at position 15 in *S. cerevisiae* and corresponding lysine at position 15 in *H. sapiens*. (B, C) Phenotypic analysis of yeast cells expressing wild type or the indicated H2A mutants. Ten-fold serial dilutions of log phase cells were spotted on plates containing rich media alone or in presence of DNA damaging agent methylmethane sulfonate (MMS). *H2A-S15E* showed a slight growth defect in presence of the drug. Whereas, both *H2A-S15A/E* in background with *H2A-S129A* showed stronger sensitivity than their respective single mutants. *H2B-T129A* did not influence the growth of *H2A-S15A/E*. (D, E) ChIP-qPCR showing γ-H2A at an inducible DSB is not affected by *H2A-S15A/E*. Cells were grown in YP-raffinose overnight till log phase followed by addition of galactose to induce DSB for 3 hours before cross-linking the chromatin. Primer pairs used flanked the DNA sequence on both left and right side at 3Kb, 7Kb and 10Kb of the DSB at *MAT* locus. Anti-H4 signal was used to normalize for histone occupancy. The data represents mean of two independent biological replicates. Error bars indicate the range between the two biological replicates. (F) *H2A-S15A/E* affects the acetylation of histone H4 and H2A. Western blot analysis of yeast whole cell extracts (WCE) with anti-H4ac reveals a slight reduction in H4ac and H2Aac in *H2A-S15A* and to a lesser extent in *H2A-S15E* compared to wild type. Histone H3 was used as loading control. (G) *H2A-S15A/E* show reduced H4ac signal in regions close to the DNA break site. Signal for H4ac was normalized for histone occupancy on total H4 signal. ChIP-qPCR was performed as described in (D), with primers flanking 150bp, 1kb, 3Kb left and 0.5Kb, 1kb, 3Kb right of DSB.

This led us to investigate whether H2AS15 had a role in the repair of DNA breaks. Our results indicate that H2AS15ph is induced by Mec1^ATR^ over a large domain of chromatin around the DSB. We also present data arguing that H2AS15ph regulates Rad9 binding and DNA end resection. Altogether, these results suggest an evolutionary conserved mechanism, similar to the mammalian H2AK15 ubiquitin/acetylation switch, modulating yeast Rad9^53BP1^ function on the chromatin surrounding DNA breaks.

## RESULTS

### H2AS15ph function is distinct from H2AS129ph in response to DNA damage

In order to investigate if H2A-S15 has a role in DNA damage response, we mutated H2A-S15 to alanine (non-phosphorylation mimicking) or glutamic acid (phosphorylation mimicking). In the presence of DNA damaging agent methyl methanesulfonate (MMS), *H2A-S15E* mutant cells show a slight growth defect, which is not seen for *H2A-S15A* (**Fig. 1B**). However, in combination with the *H2A-S129A* mutant, both *H2A-S15A/E* show stronger growth defect compared to single *H2A-S129A* mutant cells (**Fig. 1B**). Next, we tested if the *H2A-S15* mutants had any genetic interaction with the other PIKK-dependent histone phosphorylation induced in response to DSBs in yeast, H2BT129ph (10). *H2A-S15A/E H2B-T129A* double mutant cells do not show any additional growth defect compared to *H2A-S15* single mutants in presence of MMS (**Fig. 1C**). These results indicate that H2AS129 and H2AS15 phosphorylations play distinct roles in response to DNA damage induced by MMS.

To decipher if the *H2A-S15* mutants affect H2AS129 phosphorylation *in vivo*, we performed ChIP-qPCR at an inducible DSB. We used the yeast background in which a single persistent DNA double strand break is created at the *MAT* locus by the homothallic endonuclease (HO), inducible by galactose-containing media. The induced H2AS129ph signal detected on each side of the DSB in wild type (WT) cells corresponds to what was previously reported (10, 12). Both *H2A-S15A/E* mutants show similar pattern of H2AS129ph compared to WT cells, not affecting the induction or spreading of H2A-S129ph beside a very small increase in the *H2A-S15E* mutant (**Fig. 1D-E**). These results again support a role of H2AS15ph in DNA damage response/repair that is distinct from H2AS129ph.

The lysine residues of H2A and H4 N-terminal tails are known targets of the NuA4 acetyltransferase complex (30). Studies have described histone modifications modulating the occurrence of other modifications in chromatin, thus forming a network of histone crosstalk (16). H4S1ph carried out by CK2 has been shown to inhibit acetylation of various lysine residues on the N-terminal tail of H4 and play an active role in DSB repair as well as transcription (19, 20). This raised the question whether H2AS15ph could influence acetylation of lysine residues on the H2A and H4 N-terminal tails. Western blot analysis of whole cell extracts from yeast cells harboring the *H2A-S15* point mutants and grown in normal conditions reveals a small decrease in both H2A and H4 acetylation, more noticeable for the *H2A-S15A* mutant compared to wild type (**Fig. 1F**). We further investigated *in vivo* if the *H2A-S15* mutants affected the acetylation of histones around DSBs. Both *H2A-S15A/E* mutants show a decrease in H4 acetylation signal but only close to the DNA break site (**Fig. 1G**). These results suggest that H2AS15ph has a small but significant effect on other histone modifications in regions surrounding the DSB.

### H2AS15ph is induced over a large chromatin domain surrounding DNA double-strand breaks

To characterize the dynamics of H2AS15ph in response to DNA damage, we produced and purified an antibody against the *S. cerevisiae* H2A N-terminal portion with Ser15 bearing the phosphorylation mark. To demonstrate that H2AS15ph is indeed regulated by DNA damage, we performed immunoblotting with purified yeast native chromatin. As shown in **figure 2A**, H2AS15ph signal increases with MMS treatment, similarly to H2AS129ph. A control using phosphatase treatment of the chromatin confirms that the signal detected by the antibody is indeed phosphorylation-specific (**Fig. 2B**).

**Fig 2:**
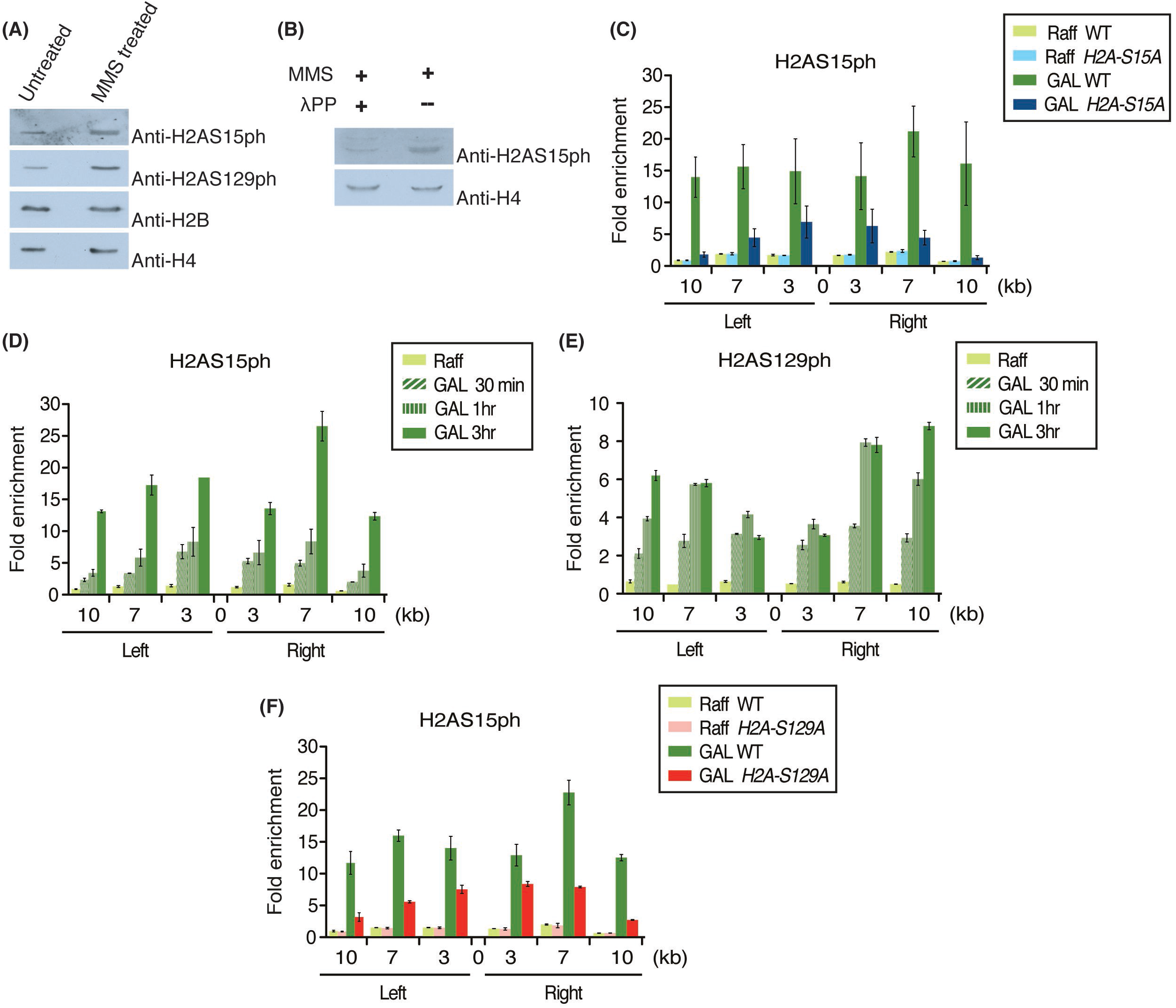
H2AS15ph is locally induced upon DNA double strand break formation and spread over several kilobases. (A) H2AS15ph signal increased when yeast cells are treated with DNA damaging agent, MMS. Western blot analysis was performed with 300ng of native chromatin purified from yeast cells treated with DMSO (untreated control) or MMS. Anti-H2AS129ph was used as positive control for the MMS treatment. Antibodies against histone H2B and H4 were used as loading control. (B) Anti-H2AS15ph is specific for recognizing a phosphorylation mark on H2A. Western blot analysis was performed with 300ng of native chromatin purified form MMS-treated yeast cells after incubation with or without lambda protein phosphatase (λPP). Lane with λPP treatment showed a significant decrease in signal for H2A-S15ph. Histone H4 was used as loading control. (C) H2AS15ph spreads on both sides of the HO endonuclease-induced single DSB at the *MAT* locus. Signal for H2AS15ph showed enrichment after the induction of the DSB in wild type but not in *H2A-S15A*. ChIP-qPCR was performed as described in Figure 1D. Fold enrichment was calculated as ratio of %IP/ input normalized on total H4 for histone occupancy at indicated loci around the DSB to the signal at the negative control locus intergenicV. (D, E) Kinetics of H2S15ph and H2AS129ph accumulation after the induction of the HO DSB. ChIP samples were analyzed after 0, 30min, 1hr and 3hrs of HO induction. Anti-H4 signal was used to normalize for histone occupancy. Enrichments were measured as described in (C). (F) Decreased H2AS15ph signals in cells lacking γH2A. ChIP-qPCR of H2AS15ph before and after HO induction in wild type and *H2A-S129A* mutant cells. Enrichments were measured as described in (C).

To determine whether H2AS15ph occurs at the DSB, we performed ChIP-qPCR at the *MAT* locus before and after HO induction. The signal detected for H2AS15ph shows a clear increase around the DSB site in WT background spanning over 10 kilobases on each side of the break (**Fig. 2C**). As a control to check the specificity of the signal detected by the antibody, the *H2A-S15A* mutant cells do not show such signal over a large domain after induction of the DSB (**Fig. 2C**). However, close to the break site, *H2A-S15A* mutant cells do show a slight increase which likely reflects some non-specific cross-reactivity of the antibody with another mark. To characterize the dynamics of H2AS15ph appearance around the DSB, the ChIP-qPCRs were performed at different time-points after induction of the HO endonuclease. Interestingly, appearance of H2AS15ph seems delayed in comparison to H2AS129ph, reaching maximum levels throughout the region after 3hrs, while H2AS129ph reaches its maximum levels at 3 and 7kb within an hour (**Fig. 2D-E**). Since H2AS129ph seems deposited before H2AS15ph, we tested if it is required for H2AS15 phosphorylation to occur. ChIP-qPCR in wild type and H2AS129A mutant cells does show a significant decrease of H2AS15ph signal around the HO-induced DSB, but the mark is still present (**Fig. 2F**). These results indicate that γH2A/H2AS129ph can influence phosphorylation of H2AS15 around the DSB but is not *per se* required for it to occur, in agreement with the increase DNA damage sensitivity of the double mutants (**Fig. 1B**). Altogether, these data clearly demonstrate that H2AS15ph is actively deposited over a large region of chromatin surrounding DNA double-strand breaks.

### Mec1^ATR^ is solely responsible for H2AS15 phosphorylation at DSBs

Tel1^ATM^ and Mec1^ATR^ signaling kinases phosphorylate histone H2AS129 to form γ-H2A (7, 11, 31). They also target various proteins to modulate DSB repair and checkpoint activation. Recruitment of Tel1 to DSB requires Xrs2 (mammalian NBS1), subunit of the Mre11-Rad50-Xrs2 (MRX) complex (32). Deletion of *XRS2* affects Tel1-dependent checkpoint signaling and telomere maintenance functions (33). Yeast Lcd1/Ddc2 (mammalian ATRIP) is responsible for the recruitment of Mec1 to RPA-coated single-stranded DNA (ssDNA) produced during end resection (34). Mec1 has been shown to be involved in spreading of γ-H2A in *trans* onto adjacent undamaged chromosomes in close physical proximity (10, 35). Deletion of *LCD1* leads to loss of Mec1 recruitment to RPA-coated ssDNA and, therefore, Mec1-dependent cell signaling in response to damage (36, 37).

*lcd1*Δ and *lcd1*Δ*xrs2*Δ mutant cells are respectively defective in the recruitment of Mec1 and both Mec1/Tel1 at the DSB. In *lcd1*Δ cells, where Tel1 recruitment and signaling is unaffected, ChIP-qPCR experiments show that the signal for γ-H2A close to the break is similar to wild type cells (**Fig. 3A**). However, farther from the break, at 7Kb and 10Kb, the signal is significantly reduced, showing its relation to Mec1. In *lcd1*Δ*xrs2*Δ cells, where both Mec1 and Tel1 recruitments are abrogated, the γ-H2A signal is completely abolished throughout the region (**Fig. 3A**). This is similar to what has been observed for *mec1*Δ and *mec1*Δ*tel1*Δ mutant cells in previous studies (10). Loss of *MEC1* affects γ-H2A spreading away from the break site, whereas loss of *MEC1* and *TEL1* results in complete loss of γ-H2A.

**Fig 3:**
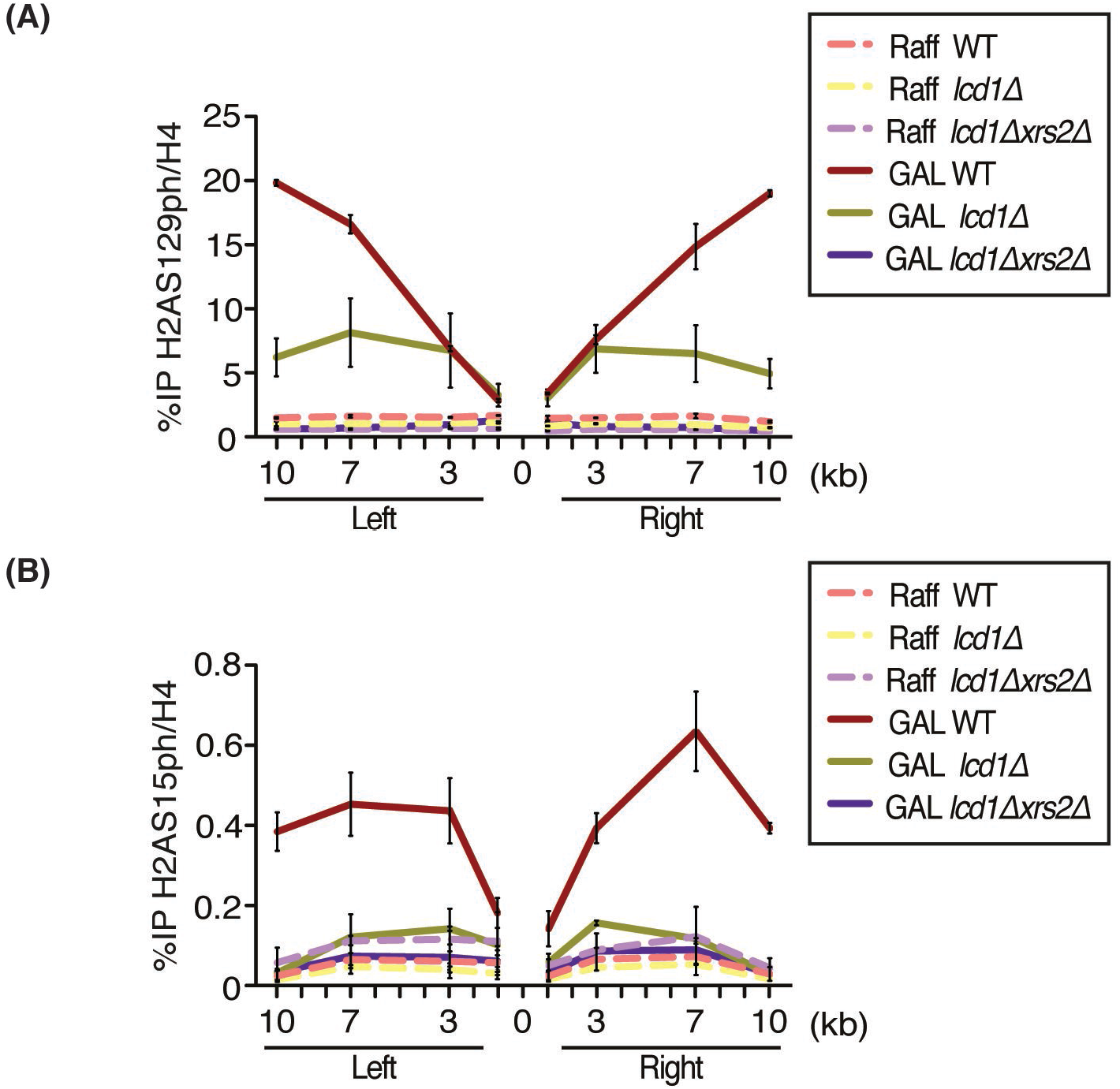
Mec1^ATR^ mediates phosphorylation of H2AS15 *in vivo*. (A) Phosphorylation of H2AS129 carried out by Mec1^ATR^ and Tel1^ATM^. Signal for H2AS129ph at DSB in *lcd1*Δ (loss of Mec1) is similar to wild type proximal to the break but much reduced distal to the break. H2AS129ph signal is abolished in *lcd1*Δ*xrs2*Δ (loss of Mec1 and Tel1). The line representing the signal for GAL *lcd1*Δ*xrs2*Δ is hidden behind since it has similar values as uninduced/raffinose samples. (B) Phosphorylation of H2AS15 is only mediated by Mec1^ATR^. Loss of H2AS15ph signal in *lcd1*Δ and *lcd1*Δ*xrs2*Δ compared to WT. All yeast stains used in the above experiments have *sml1*Δ background. ChIP-qPCR was performed as described in Figure 1D, primer pairs flanking 1Kb left and right of DSB were also used.

A large scale phosphoproteomic study has detected phosphorylation of H2AS15 and linked it to Mec1 (29). To investigate this, we carried out ChIP-qPCR using the H2AS15ph antibody to study its spreading around the DSB in *lcd1*Δ and *lcd1*Δ*xrs2*Δ mutant cells. As shown above, wild type cells show enrichment for H2AS15ph over a large region on each side of the DSB (**Fig. 3B**). However, *lcd1*Δ as well as *lcd1*Δ*xrs2*Δ mutant cells do not show any H2AS15ph signal. These results demonstrate that, unlike γ-H2A which is deposited by both the Mec1 and Tel1 kinases, H2AS15 phosphorylation is carried out solely by Mec1, which links it to DNA end resection after DSB formation.

### H2AS15 is required to maintain fidelity of resection

Previous studies have shown that the *H2A-S129A* non-phosphorylation mutant leads to faster resection from the HO-induced DSB compared to wild type cells (38, 39). This increased resection may be attributed to the inability of Rad9 to bind H2AS129ph through its BRCT domain, an essential interaction for its recruitment on chromatin (17, 27, 28, 40, 41). Budding yeast Rad9, like its mammalian homolog 53BP1, functions in slowing down or keeping a check on the rate of DNA end resection (42-45). To investigate whether H2AS15ph may affect resection, we used an antibody against Rfa1, RPA largest subunit, as a proxy for the ssDNA generated by resection, in ChIP-qPCR at the HO-induced DSB. Interestingly, *H2A-S15E* (phosphorylation mimicking) mutant cells show increased RPA binding near the break, whereas *H2A-S15A* mutant cells behave like wild type (**Fig. 4A**). The *H2A-S129A* cells also show increased RPA signals, reflecting higher ssDNA/resection in this mutant. Double mutants *H2A-S15A/E H2A-S129A* have similar increased RPA signals as *H2A-S129A* cells, maybe slightly less than *H2AS15E* cells at some locations (**Fig. 4A**). To confirm that these RPA signals are the consequence of increased DNA end resection, we directly measured by qPCR the DNA signal 5kb away from the DSB, compared to an unrelated genomic locus. In this assay, a very significant and similar decrease of signal is detected in *H2A-S15E, H2A-S129A* and double mutants, but not in *H2A-S15A* (**Fig. 4B**). These results indicate that H2AS15 phosphorylation leads to increased resection at DSBs, similar to the effect of losing H2AS129 phosphorylation. Furthermore, the effects of each mutant are non-additive, suggesting that they are mechanistically related. An interesting possibility is whether H2AS15ph is also recognized by the BRCT domain of Rad9, this domain being well characterized for its ability to bind H2AS129ph (27). Surprisingly, *in vitro* peptide binding assays with recombinant Rad9 indicate a clear binding preference for the histone H2A tail peptide, but in its non-phosphorylated form (**Fig. 4B**). Importantly, this binding specificity is also confirmed using yeast whole cell extracts in peptide pull-downs and detecting endogenous Rad9 by immunoblotting (**Fig. 4C**). In this case, inducing DNA damage in the cells does not affect endogenous Rad9 ability to bind the H2A peptide *in vitro*. This ability of H2AS15ph to block Rad9 binding to the H2A histone tail may in part explain the increased end resection detected in *H2A-S15E* mutant cells but not in *H2A-S15A* (**Fig. 4A-B**). As these results suggest that Rad9 may directly bind histone H2A tail in chromatin *in vivo*, and that H2AS15 phosphorylation may regulate this interaction, we analyze Rad9 binding near the HO-induced DSB by ChIP-qPCR. In wild-type conditions in this specific yeast genetic background, we could only detect Rad9-myc signal by ectopic expression from a plasmid. Furthermore, the specific signal based on the empty vector control could only be detected very close (150bp) on each side of the HO break (**Fig. 4E**). At these locations, Rad9 binding is indeed abrogated in the *H2A-S129A* mutant, as expected, and validating the specificity of the signal. In contrast, Rad9 binding does not seem affected by *H2A-S15A/E* mutations. Since H2AS15 phosphorylation occurs more downstream away from the DNA break, the Rad9 signal detected here close to the HO site may not reflect the population regulated by H2AS15ph (see (42, 46) for models of two populations/functions of Rad9/53BP1 near DNA ends versus larger domains). Another possibility is that H2AS15ph is not required for Rad9 recruitment but regulates the way it binds nucleosomes and influences end resection. Altogether, these results implicate the DNA damage-induced H2AS15ph histone mark in the regulation of end resection at DSBs.

**Fig 4:**
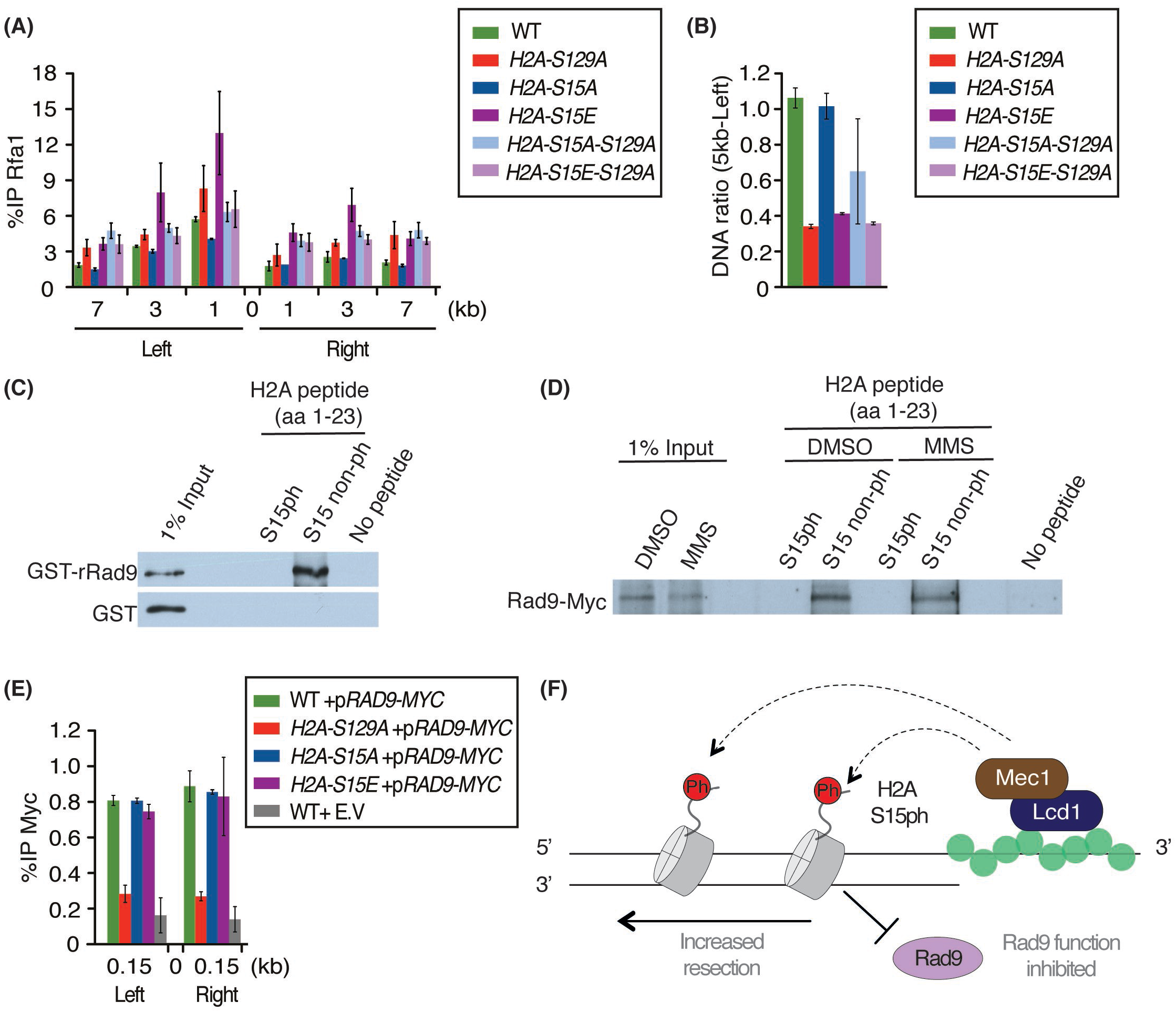
H2AS15 regulates resection of DNA ends at DSB and Rad9^53BP1^ binding *in vitro*. (A) *H2A-S15E* mutant has increased DNA end resection compared to *H2A-S15A* and wild type backgrounds, as shown by single-strand DNA binding RPA signal. *H2A-S129A* is used as a positive control as it also leads to higher resection/RPA binding. ChIP-qPCR was performed as described in Figure 1D with antibody against Rfa1 (largest RPA subunit) and primer pairs flanking 1Kb, 3Kb and 7Kb left and right of DSB. (B) Increased DNA end resection in the *H2A-S15E* mutant is also confirmed by directly measuring DNA signal loss near the HO DSB. After 3hrs in inducing media, qPCR was used to measure DNA at 4.8kb Left of the HO cut-site and at control locus in the genome (*PRE1*). Signal ratios on the control locus show are decreased in *H2A-S15E* and *H2A-S129A* mutants, indicating similar higher end resection in these cells. (C) Peptide pull-down assay with GST-tagged recombinant Rad9 and indicated H2A N-terminal peptides. rRad9 binds to unmodified peptide over phosphorylated peptide. Empty GST is used as negative control. (D) Peptide pull-down assay as in (B) but using extracts from yeast cells treated or not with MMS and expressing myc-tagged Rad9. (E) Binding of Rad9 close to the HO DSB is not affected by the H2AS15 mutants *in vivo*. ChIP-qPCRs of Rad9-Myc ectopically expressed in the indicated backgrounds show significant binding within 150bp of the break, which is lost in the *H2A-S129A* mutant but not in *H2A-S15E*. An strain with an empty vector (E.V.) is used as control for background non-specific Myc signal. (F) Model for the role H2AS15ph in DSB repair. Mec1 phosphorylates H2AS15 in the region surrounding the DSB. This phosphorylation blocks Rad9 binding to the H2A tail in the bound nucleosomes, which disrupts its function and results in increased resection. H2AS15ph may function as a dynamic switch to modulate resection, phosphorylation enabling resection and non-phosphorylation preventing hyper-resection.

## DISCUSSION

Histone tails are sites of various post-translational modifications (PTMs) such as phosphorylation, acetylation, methylation, ubiquitylation and SUMOylation. These modifications are recognized by reader proteins containing specialized domains such as ubiquitin binding domains, BRCT and FHA domains that bind phosphorylation, Chromodomains, MBT domains, Bromodomains and PHD fingers that recognize methylation or acetylation. Proteins and protein complexes use these interactions for their proper recruitment, retention and/or orientation at the target sites to carry out their respective functions (47, 48).

In this study, we characterized yeast H2AS15ph as a novel histone modification in the context of DNA damage. This modification is induced in the presence of DNA damaging agent MMS, which is a known alkylating agent and produces DSBs as cells replicate their DNA in S-phase. A large-scale proteomic study had identified H2AS15ph by mass spectrometry and implicated Mec1^ATR^ for its detection (29). Using *in vivo* analysis, we show that indeed Mec1, and not Tel1, is responsible for the induction of this mark around DNA breaks, unlike γ-H2A (H2AS129ph) which is deposited by either Tel1 or Mec1 (**Fig. 3**). While the spreading of γ-H2A at DSBs is not affected in *H2A-S15A/E* mutant cells, the *H2A-S129A* mutant does affect the levels of H2AS15ph (**Fig. 1C-D, 2F**). Furthermore, a decrease of chromatin acetylation is detected in *H2A-S15A/E* mutant cells near the DNA break (**Fig. 1F-G**). We know from previous studies that acetylation of histone H4 and H2A by the NuA4 complex plays an important role for the recruitment of various chromatin remodelers like INO80, RSC and SWI/ SNF required for repair by HR (12, 49-51). Future work will be needed to determine if H2AS15 affects the recruitment of these remodelers.

A most interesting finding reported here argues that yeast H2AS15ph corresponds to an evolutionary conserved regulatory function to regulate 53BP1/Rad9 binding to the H2A tail and DNA end resection. The yeast H2AS15 residue seems the functional homolog of mammalian H2AK15 in modulating resection which in turn affects DNA repair pathway selection, albeit through different PTMs. In mammals, when 53BP1 binds H2AK15ub in the vicinity of DSBs, it inhibits resection and favors the NHEJ pathway. This binding inhibits TIP60-mediated acetylation of H4 and H2A tails in chromatin, including H2AK15, which otherwise would block ubiquitylation by RNF168 and 53BP1 binding, favoring resection and HR (25). *H2A-S15E* mutant cells show increased resection, as seen with *H2A-S129A* mutant cells where Rad9^53BP1^ is incapable to bind and inhibit or slow down resection (**Fig. 4A-B**). *In vitro* data suggest that Rad9 binds the H2A N-terminal tail in its unmodified state and this binding is abolished when H2AS15 is phosphorylated (**Fig. 4C-D**). On the other hand, this regulated interaction does not seem to affect Rad9 recruitment to the DSB (**Fig. 4E**). Since Rad9 is recruited to nucleosomes near DSBs though bivalent interaction with H2AS129ph and H3K79me, our results suggest that Rad9 interaction with the H2A tail is not required for its recruitment but can regulate its function linked to DNA end resection. 53BP1 also interacts with nucleosomes through three interfaces, i.e. H2AK15ub-UDR, H4K20me-Tudor and H2A.XS139ph-BRCT. But, as in yeast, only two are required for its binding to nucleosomes, γH2AX-BRCT being dispensable (24, 46, 52). We proposed a model depicted in (Fig. 4F) where resection of DNA ends produces ssDNA which gets coated by RPA and recruits Mec1^ATR^ through its binding partner Lcd1/Ddc2^ATRIP^ (34, 36). Mec1 then phosphorylates H2AS15 which inhibits Rad9 function and therefore increases resection. H2AS15ph may be a dynamic mark, switching between phosphorylation and non-phosphorylation states to fine-tune the process of resection. This conserved function of the H2A tail to regulate Rad9/53BP1 binding and resection is even further supported by a recent report showing phosphorylation of the ubiquitin moiety of H2AK15ub (H2AK15pUbT12) to inhibit 53BP1 binding and favor resection (53).

Future work will address whether specific proteins may bind H2AS15ph. While our peptide pull-downs with yeast extracts followed by mass spectrometry did not yield clear results (data not shown), we speculate that other DNA repair proteins such as Rtt107 and Dpb11 which have BRCT domains could recognize H2AS15ph. Rtt107 was recently shown to bind H4T80ph and γ-H2A to counteract Rad9-modulated checkpoint activity (54). Dpb11 has been shown to act as a scaffold protein for various checkpoint proteins and is involved in Rad9 recruitment/retention at DSBs (55, 56). H2AS15 may also act redundantly with other DNA damage induced histone phosphorylation marks and function as a scaffold for proper orientation or retention of repair factors.

## MATERIAL AND METHODS

### Yeast strains

All yeast strains used in the current study are from FY406 (57) and JKM179 (58) (*hta1-htb1Δ:*:*NatMX, hta2-htb2Δ::KANMX*) backgrounds with deletion of both copies of H2A and H2B genes. These strains carry an ectopic copy of *HTA1-HTB1* on low copy vector pRS413 (in FY406) or pRS414 (in JKM179). Gene deletion and point mutations were obtained through standard PCR techniques. Plasmid shuffling using 5-Fluoroorotic acid (5-FOA) was used to introduce the plasmid with point mutations in respective backgrounds. Yeast background JKM139 (58) with deletion of *sml1Δ*::*NATMX, lcd1Δ*::*KANMX, xrs2Δ*::*TRP1* was used in this study. QY362 (JKM139 *RAD9-13MYC::KANMX*) was used for peptide pull down experiments while the Rad9-myc ChIP-qPCR were done in the JKM179 background with *HTA1/2-HTB1/2* deletions covered wild type or mutant H2A and ectopic Rad9-Myc expression from a pRS315 vector (40). General lithium acetate method was used to transform the yeast cells. Cells were grown in YPD (1% yeast extract, 2% peptone, 2% dextrose) at 30°C until early log phase. For spot assay 1:10 dilution of the culture was spotted and grown at 30°C for 3-4 days.

### Chromatin Immunoprecipitation (ChIP)

ChIP-qPCR was performed as described previously (59). In brief, cells were grown overnight in YP-Raffinose until OD_600_ of 0.5-1, at this point galactose (2%) was added to induced DSB at *MAT* locus for 3 hours. This was followed by cross-linking of cells with formaldehyde and lysing with bead-beater. Sonication was performed on Diagenode Bioruptor to obtain chromatin size of ∼200-500bp. 100μg of chromatin was used to setup the immunoprecipitation reaction with anti-H2A-S129ph (Upstate 07-745), anti-H2A-S15Ph (antibody was produced in rabbit using the peptide sequence: Ac-GSAAKA(pS)QSRSAK-C. Affinity purification was performed to improve the specificity of the antibody, with phosphorylated versus non-phosphorylated peptides. Antibody was produced by Thermo Fisher Scientific), anti-Rfa1 (Agrisera AS07214), anti-H4ac (Upstate 06-946), anti-H4 (Abcam 7311) and anti-Myc (9E10 Babco MMS150R) and reaction was allowed to mix on a wheel overnight at 4°C. Next morning Protein-A agarose beads (protein-G magnetic beads for anti-Myc) were added followed by additional 3-4 hours on a wheel at 4°C. After multiple washing steps the DNA was eluted and incubated at 65°C overnight for de-cross linking followed by phenol-chloroform extraction of DNA. This DNA was resuspended in NTE and used for qPCR. LC480 LightCycler (Roche) was used to quantify DNA with the primer pairs as mentioned in the respective figures to calculate % of immunoprecipitation (IP)/ input. The data represents mean of two independent biological replicates. Error bars indicate the ranges between the two biological replicates. Fold enrichment represents ratio of %IP/input at indicated loci around DSB normalized on signal at negative-control locus IntergenicV. Levels of HO-induced DNA break was verified by qPCR across the HO cut-site (normalized on control locus) and were similar in all experiments. Direct measurement of end resection by qPCR was done using primers 4.8kb left of the HO cut-site and presented as a ratio of signal over a control locus (*PRE1*, (60)). Primer sequences are available upon request.

### Purification of native yeast chromatin

Native yeast chromatin was prepared as described previously (61), cells were treated with MMS (0.05%) or DMSO (control) for 2 hours to induce DNA damage. Lambda protein phosphatase (λPP) (NEB) was used to dephosphorylate the purified chromatin. The reaction mixture was incubated for 30 mins at 30°C followed by booster of λPP and additional incubation for 30 mins at 30°C.

### Recombinant protein purification

Recombinant Rad9 (C-terminal containing Tudor and BRCT domains) protein was purified as described previously (62). In short, bacterial cells were grown overnight at 16°C with IPTG induction. Next day cells were lysed with lysozyme followed by sonication. The soluble portion was incubated with glutathione sepharose (GE healthcare) beads for 3-4 hours at 4°C and eluted with glutathione. The eluted protein concentration was quantified by running on SDS-PAGE with known BSA standards followed by coomassie staining.

### Peptide-pull down

Peptide-pull down was performed with synthetic peptide corresponding to H2A N-terminal (aa 1-23) obtained from AnaSpec. 1μg of peptide was incubated with 1μg of GST-tagged protein overnight on wheel at 4°C in binding buffer (50 mM Tris–HCl pH 7.5, 50 mM NaCl, 0.1% NP-40, 1 mM PMSF and phosphatase inhibitors). Next morning streptavidin dynabeads were added to the reaction mixture and kept at 4°C for two additional hours followed by washing, running on SDS-PAGE and western blotting with anti-GST (Sigma G1160).

Peptide-pull downs with yeast cell extracts were performed by growing cells in YPD and treating with MMS (0.05%) or DMSO for 2 hours (final OD_600_ ∼1.0-1.5). Cells were centrifuged, the pellet was washed with cold milliQ water, resuspended in sensitization buffer (100mM Tris-HCl pH 9.4, 10mM DTT) and incubated at 30°C on a wheel for 15 mins. It was then centrifuged at 500g for 5mins at 4°C, resuspended in 30ml of spheroplasting buffer (1.2M Sorbitol 20mM Hepes pH 7.5), centrifuged again, resuspended in 40ml of spheroplasting buffer with lyticase and incubated on wheel at 30°C for 90 mins. After centrifugation at 500g for 5 mins, wash with wash buffer (1.2M Sorbitol, 20mM Pipes pH 6.4, 1mM MgCl_2_), it was resuspended in 25ml of nuclear isolation buffer; NIB (250mM Sucrose, 60mM KCl, 1mM PMSF, 14mM NaCl, 0.8% Triton X-100, 5mM MgCl_2_, 1mM CaCl_2_, 15mM MES), mixed well to remove clumps and kept on ice for 20 mins. After centrifugation and two washes with NIB (until white nuclear pellet was obtained), the pellet was resuspended in pull-down buffer (400mM NaCl, 1mM PMSF, 50mM Tris-HCl pH8, 0.5% NP-40, 5 mM β-glycerophosphate, 10 mM sodium butyrate, 5 mM NaF) to get 300mM NaCl in the final volume, followed by incubation on rotating plate at 4°C for 20 mins. After centrifugation at 14,000rpm at 4°C for 30 mins, the supernatant was collected (nuclear extract). Biotinylated peptides were bound to streptavidin dynabeads in peptide binding buffer (150mM NaCl, 50mM Tris-HCl pH8.0, 0.1% NP-40, 5 mM β-glycerophosphate, 10 mM sodium butyrate, 5 mM NaF) by incubating at room temperature on wheel for 30 mins. Peptide-dynabeads were washed with protein binding buffer (140mM NaCl, 50mM Tris-HCl pH 8.0, 0.1% NP-40, 0.5mM DTT, 10μM ZnCl_2_, 5 mM β-glycerophosphate, 10 mM sodium butyrate, 5 mM NaF) followed by addition of 600μg of nuclear extract to each pull-down reaction and incubated at 4°C on wheel for 2 hours. Nuclear extract that was incubated with phospho peptide was initially incubated with non-phospho peptide, on the other hand, nuclear extract for non-phospho peptide was initially incubated with phospho peptide for 2 hours at 4°C on wheel. After 3-4 washed with final wash buffer (350mM NaCl, 50mM Tris-HCl pH8.0, 0.1% NP-40, 0.5mM DTT, 10μM ZnCl_2_, 5 mM β-glycerophosphate, 10 mM sodium butyrate, 5 mM NaF), Laemmli buffer was added, boiled and resolved in SDS-PAGE followed by western blotting with anti-Myc (9E10 Babco MMS150R).

## Acknowledgments

We thank Xue Cheng for important advice. We are grateful to Jim Haber for sharing reagents/strains and stimulating discussions. This work was supported by a grant from the Canadian Institutes of Health Research (CIHR) to J.C. (FDN-143314). S.A. held a Desjardins/Fondation du CHU de Québec studentship. J.C. holds the Canada Research Chair in Chromatin Biology and Molecular Epigenetics.

